# The impact of RNA chemical probing reagents on RNA binding proteins

**DOI:** 10.1101/2024.03.21.586119

**Authors:** David Klingler, Lucy Fallon, Daniel Cohn, Amy Crawford, Liza Marcus, Alisha N Jones

## Abstract

RNA chemical probing experiments are a broadly used method for revealing the structure of RNA, as well as for identifying protein binding sites. This is beneficial for expanding our understanding of biological processes governed by protein-RNA complex interactions, as well as facilitating the identification of complex inhibiting molecules. The reagents commonly used in chemical probing experiments are highly reactive, methylating or acylating flexible RNA nucleotides. The highly reactive nature of the chemical probes means that they can also react with nucleophilic amino acid side chains, and subsequently affect protein-RNA binding events. We combine molecular dynamics (MD) simulations, matrix-assisted laser desorption ionization mass spectrometry (MALDI-MS), and nuclear magnetic resonance (NMR) experiments to show that commonly used RNA chemical probes react with protein amino acids, and demonstrate that this effect alters protein-RNA binding interactions through binding shift assays. We discuss the implications of this phenomenon in elucidating the protein-RNA interaction interface using chemical probing experiments.

**GRAPHICAL ABSTRACT:** 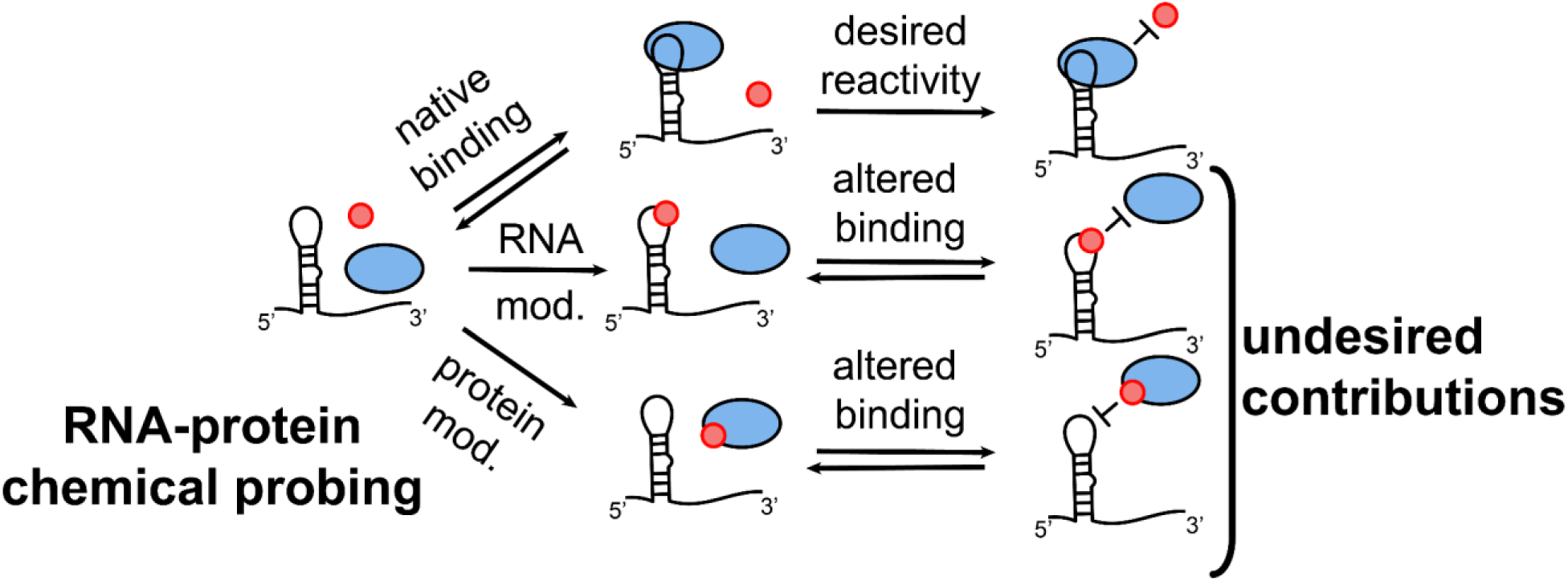

## INTRODUCTION

RNA structure is best described as an ensemble of various conformations (1). The dynamic nature of RNA makes experimentally characterizing its structure challenging. Biochemical experiments such as selective 2’-hydroxyl acylation analyzed by primer extension (SHAPE), and dimethyl sulfate (DMS) chemical probing are used to determine the secondary structure of conformationally dynamic RNA molecules, as well as to identify tertiary structural contacts of RNAs both *in vitro* and in cells (2–6). This method has been used to elucidate the secondary structure of both small and large RNA molecules, from micro-RNAs to entire viral genomes (7, 8). Further, chemical probing is often used to identify protein and other ligand binding sites on RNA molecules: regions of the RNA that demonstrate different chemical reactivities in the presence of a binding partner versus in the absence of it are typically interpreted as binding sites (9–11).

The chemical probes used in SHAPE experiments react rapidly with the 2’-hydroxyl group of the ribose sugar of unpaired or everted nucleotides, covalently modifying the ribose (12). DMS functions similarly: it binds and methylates the N1 position of adenine, and the N3 position of cytosine (13). At higher pH, uridines and guanosines are also susceptible to DMS methylation (14). The covalently modified RNA is then sequenced, using either reverse transcription-stop (RT-stop) or next-generation sequencing (RT-map) methods, which permits identification of the nucleotides that reacted with the chemical probes (7, 13). The reactivities are then used to improve the prediction of RNA structure with *in silico* folding approaches.

One of the benefits of chemical probing experiments is that they can be used to identify sites of protein and other ligand binding in an RNA. For example, in a recent study, SHAPE was used to identify where two RNA binding proteins (RBPs), hnRNP U and hnRNP L, bound to the precursor messenger RNA MALT1 to regulate how it is spliced (15). In another study, DMS probing was used to identify how proteins bind to increase the flexibility of the 7SK noncoding RNA, thereby increasing the accessibility of neighboring binding sites (16). Chemical probing has also been used to assess ligand binding affinity when carried out at variable ligand concentrations (17–19).

These experiments have enabled researchers to better understand a plethora of biological processes that rely on ligand-RNA interactions. One aspect of using chemical probing to map protein/ligand interactions that has received little attention is the ability of the chemical probes to interact with the ligand, especially given the on-off rates of the RNA-ligand complex and the rate at which the chemical probes interact. Relative to an RNA-only control (**Figure 1A**), when a protein ligand is present, four scenarios can occur (**Figure 1B-E**). First, binding of the protein to the RNA can prevent the probe from interacting with the RNA molecule (**Figure 1B**). In the second scenario, the probe is bound to the RNA, and prevents the recognition of the RNA by the protein (**Figure 1C**). Most poorly understood is the ability of the chemical probes to interact with the protein (and/or the RNA), impeding complex formation (**Figure 1D, E**). Lastly, combinations of these scenarios may occur. It is important to note that if chemical probing is to be used to investigate binding affinities only one type of interaction, scenario B, is desired.

**Figure 1.**
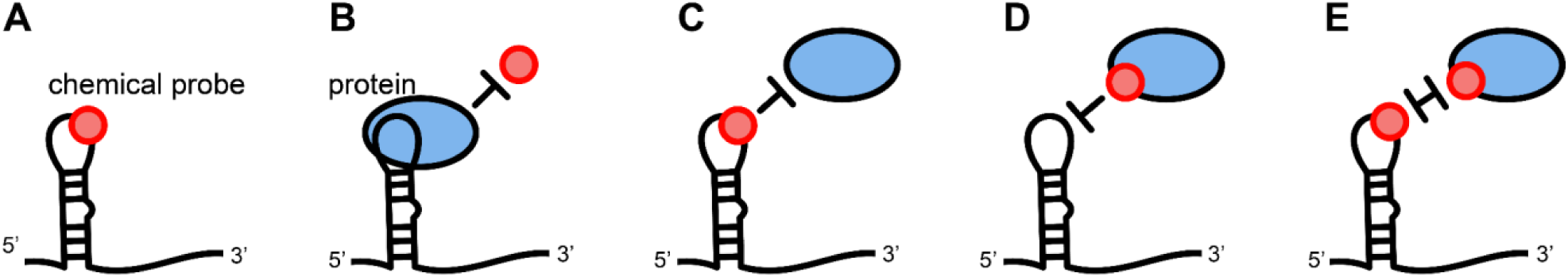
Mechanisms of chemical probing experiments. **A)** Chemical probes modify flexible or single-stranded nucleotides. When protein is present in solution, **B)** the protein can prevent binding by the probe, **C)** the probe can prevent binding of the protein, **D)** the probe can bind to the protein to prevent an interaction, and **E)** the interaction site can be doubly modified, preventing an interaction.

The lack of data for the interactions between reagents used for RNA chemical probing and proteins is remarkable considering the same reactive moieties forming the basis for RNA probing reagents have been employed to react with proteins repeatedly in the past. The isatoic anhydride scaffold has been used extensively to covalently tag proteins for the purpose of fluorescence spectroscopy (20), biotinylation (21, 22), introduction of chemical handles for further reactions (23), and gel pre-labeling (24, 25). Initial studies had suggested that the nucleobase modifying DMS was suitable for use in the presence of proteins and only *N*-Cyclohexyl-*N′*-(2-morpholinoethyl)carbodiimide methyl-*p*-toluenesulfonate (CMCT) needs be avoided due to side reactions (26). However, a more recent study employed DMS-induced methylations to establish protein-protein interaction footprints, highlighting the ability of DMS to modify proteins as well as RNA (27).

Several proteinogenic amino acids possess nucleophilic functional groups in their side chains (**Figure 2 and S1**). Some of these are known to be post-translationally modified by acylations (28) and methylations (29, 30). The chemical probing reagents are capable of reacting with side chain nucleophiles in a similar manner: isatoic anhydride and its derivatives have been reported to react with lysine (Lys) side chains (20, 21, 23). However, in line with the typical reactivity of the isatoic anhydride scaffold, chemical probing reagents like 1M7 and NMIA may also acylate the side chains of cysteine (Cys) as well as the hydroxyl groups of tyrosine (Tyr), serine (Ser), and threonine (Thr) (**Figure 2A**) (31). The products of acylations of cysteine would be thioesters, which are known to be unstable. However, some thioesters, such as acetyl-CoA, are found in the cellular context. DMS-induced methylations have been reported for Lys, histidine (His), and glutamate (Glu) residues (27). By extension, Cys and aspartate (Asp) might also be susceptible (**Figure 2B**). However, identifying potential modification sites and reactivities for these reagents is substantially more complex in a protein than it is for a peptide or monomer. The nucleophilicity of a side chain in the context of a protein is determined not just by the type of amino acid itself: neighboring amino acid residues can significantly alter a side chain’s nucleophilicity. The catalytic triad is a typical example of an architecture enhancing the nucleophilicity of a specific residue (32). Accessibility for reagents is another major factor to consider when dealing with side chain modifications. Thus, the three dimensional structure of a protein will have an influence on the reactivity of a specific residue when presented with a modifying reagent. In this sense, proteins may behave similarly to nucleic acids when presented with chemical probing reagents.

**Figure 2.**
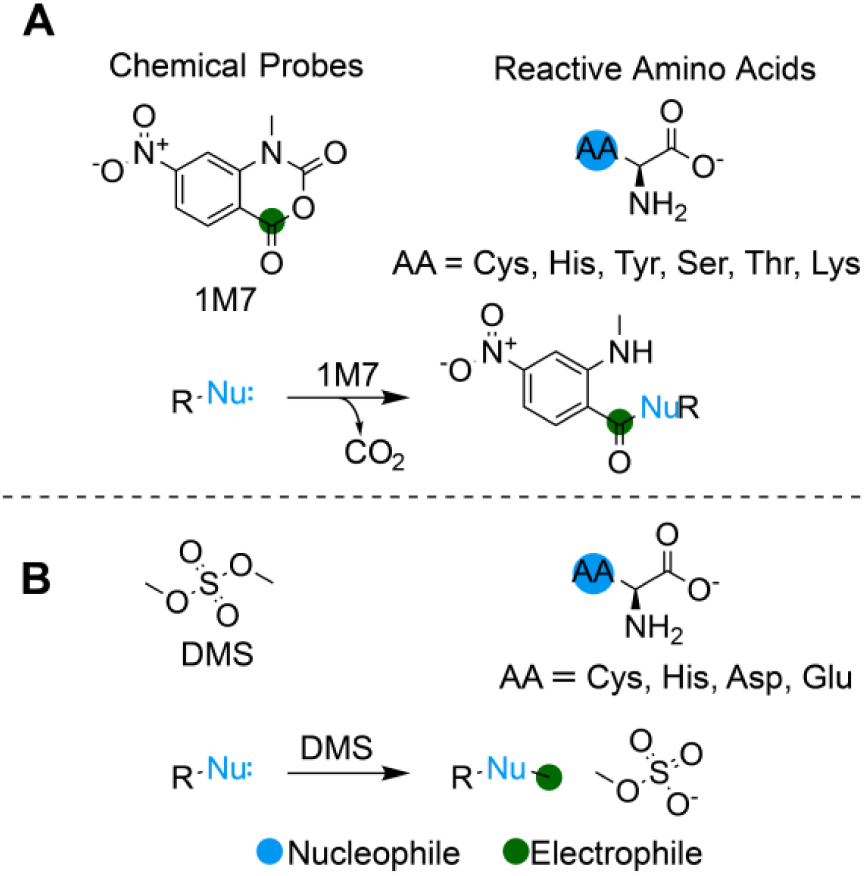
Nucleophilic amino acids that react with RNA-modifying probes. **A)** 1M7 acylates Cys, His, Tyr, Ser, Thr and Lys residues. **B)** DMS methylates Cys, His, Asp, and Glu residues.

In this study we combine binding shift assays, NMR, MALDI-MS, and MD simulations to demonstrate that the chemical probes frequently used in RNA chemical probing experiments can bind to and react with nucleophilic amino acids that reside on the surface of proteins. Our results show that the covalent modification of amino acids by the chemical probing reagents alters the protein binding surface, and can obscure the accurate deduction of protein-RNA interactions. We discuss the implications of the chemical probes reacting with amino acids in interpreting protein-RNA binding events, thus highlighting the importance of further validating ligand-RNA interactions with alternative methods following chemical probing experiments.

## RESULTS

### MALDI-MS reveals covalent modification of proteins

We used MALDI-TOF (matrix-assisted laser desorption and ionization-time of flight) mass spectrometry to determine if commonly used RNA chemical probing reagents could covalently modify proteins. MALDI-MS is perfectly suited for facile determination of intact protein molecular weights. As a model system, we used the KH3 domain of the heterogeneous nuclear ribonucleoprotein K (hnRNP K). hnRNP K possesses three RNA binding KH domains, and has several cellular roles, including regulation of dosage compensation through an interaction with XIST, and regulation of mRNA splicing (33, 34).

We treated protein solutions of 50-200 μM with a final concentration of 18 mM 1M7 (1-methyl-7-nitro isatoic anhydride), NMIA (N-methyl isatoic anhydride), BzCN (benzoyl cyanide), 2A3 ((2-Aminopyridin-3-yl)(1*H*-imidazol-1-yl)methanone), and 20 mM DMS, and compared the molecular weight with unmodified protein. These concentrations mimic the concentrations used in chemical probing experiments. The resulting mass spectra (**Figure 3A**) clearly show multiple species with increased molecular weights for hnRNP K KH3 modified by 1M7, NMIA, and 2A3 corresponding to a complex mixture of singly, doubly and, multiply modified proteins. The observed increase in molecular weight is not a result of non-covalent adduct formation since adding the 1M7 hydrolysis product to hnRNP K KH3 did not produce the same pattern in the mass spectrum (**Figure S2**). The overall modification rate is substantial given that the unmodified protein accounts only for a minor fraction in the mass spectra. We observed that 2A3 is a particularly powerful acylation reagent for the protein: while the unmodified protein still accounts for a major fraction for the other probes, it is almost undetectable for 18 mM 2A3 (**Figure 3A**, bottom). At higher concentrations of 2A3, MALDI-MS detects even higher modification rates (**Figure 3B**). In stark contrast, the observed m/z is not changed significantly by DMS and BzCN, indicating that these probes may not modify KH3 side chains (**Figure 3A**). The reactivity of DMS, in the context of RNA chemical probing, is dependent on the pH of the reaction. At pH 8.0, all four nucleotides are susceptible to DMS (14). This inspired us to repeat the DMS reaction at pH 8.0 and compare it to a control conducted at pH 6.5 by MALDI-TOF (**Figure S3**). However, the results were the same as at pH 6.5. This was surprising since DMS induced methylations have been observed in literature for other proteins (27).

**Figure 3.**
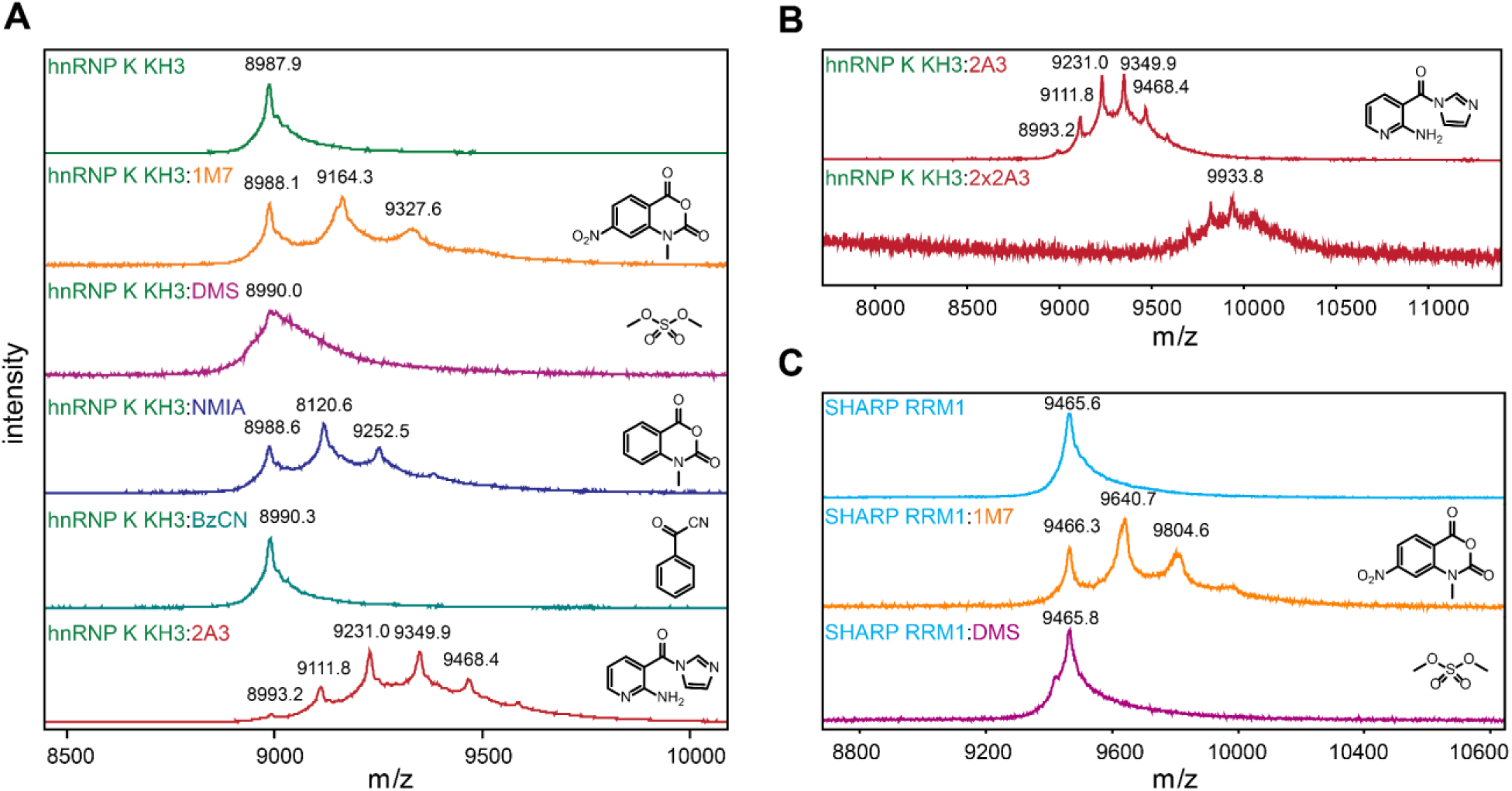
MALDI MS of proteins treated with RNA chemical probing reagents. A) hnRNP K KH3 (top) is covalently modified by 1M7, NMIA, and 2A3. Single, double and multiple modifications are observed. Treatment with DMS and BzCN did not cause modification patterns. B) 2A3 is especially potent for protein modification. Increasing the reagent concentration from 18 mM (top) to 36 mM (bottom) leads to heavily modified protein species with the unmodified variant no longer observed. C) Protein modification by RNA chemical probing reagents is also observed in SHARP RRM1. DMS and 1M7-induced changes in the mass spectra are comparable to hnRNP K KH3.

To ensure the observed covalent modifications are not a specific phenomenon observed in hnRNP K KH3, we treated the RNA recognition motif 1 (RRM1) domain of SMRT/HDAC1 Associated Repressor Protein (SHARP) with 1M7 and DMS as examples for nucleobase and 2’OH modifying reagents for comparison (**Figure 3C**). SHARP possesses four RNA recognition motifs (RRMs) that have been demonstrated to interact with several RNAs including Xist and SRA to regulate gene expression (35–37). Again, 1M7 caused covalent modification of the protein, while DMS did not cause a significant increase in molecular weight.

### MD simulations reveal 1M7 and DMS occupancy near nucleophilic amino acids

We turned to MD simulations to investigate site selectivity of the covalent modification of the proteins. In addition, we aimed to understand why we did not observe clear signs for DMS induced methylations. The simulations allow us to dissect the interaction between the chemical probing reagents and the protein, since the initial binding event of the chemical probe cannot be captured by experiments due to the subsequent reaction. As an example of a common nucleobase and 2’-OH modifying reagent, we picked the interaction of 1M7 and DMS with both RRM1 of SHARP, and the KH3 domain of hnRNP K. Both proteins possess surface and buried amino acids that could potentially interact with 1M7 and DMS.

Three 1.0 μs MD simulations of the RRM1 domain of SHARP, or the KH3 domain of hnRNP K, were performed with either 1M7 or DMS, to give a total of four different systems (SHARP + 1M7; SHARP + DMS; hnRNP K + 1M7; hnRNP K + DMS). To ensure the protein structures were stable in the simulations with 1M7 or DMS, we monitored the root mean square fluctuation (RMSF) of the protein backbone (**Figure S4**). The proteins do not deviate significantly from the reference structures in our simulations, and exhibit the greatest flexibility in the N- and C-terminal residues. The simulations of SHARP RRM1 exhibit additional flexibility in the loop region, connecting ꞵ- strands 2 and 3 (residues P39 - A48 in **Figure S4B**). We conclude that both SHARP RRM1 and the KH3 domain of hnRNP K are stable in our MD simulations.

To understand which amino acids the 1M7 or DMS molecules might associate with on the two proteins – and by extension, which amino acids may be accessible for covalent modification by these chemical probes – we monitored the occupancy of the two chemical probes around the proteins throughout the MD simulations. We calculated the radial distribution function (RDF) of 1M7 or DMS against potentially reactive amino acids, which measures the population of 1M7 or DMS at each distance from the designated amino acids. For both SHARP RRM1 and the KH3 domain of hnRNP K, the 1M7 molecules preferentially associate with aromatic residues (Phe, Tyr, Trp, His) on the protein surfaces. In the simulations of SHARP RRM1 with 1M7, the 1M7 molecules form aromatic stacking interactions with His7, Trp9, His63, and Tyr78 (**Figure 4A**), which are all potentially reactive towards 1M7 and may facilitate a reaction *in vitro*. In the MD simulations of hnRNP K KH3 with 1M7, the 1M7 molecules localize near Tyr75 and Tyr84, as well as solvent-exposed residues on helix-3 such as Gln71 and Leu76 – which brings 1M7 molecules adjacent to Glu42 and Ser80 (**Figure 4B**).

**Figure 4.**
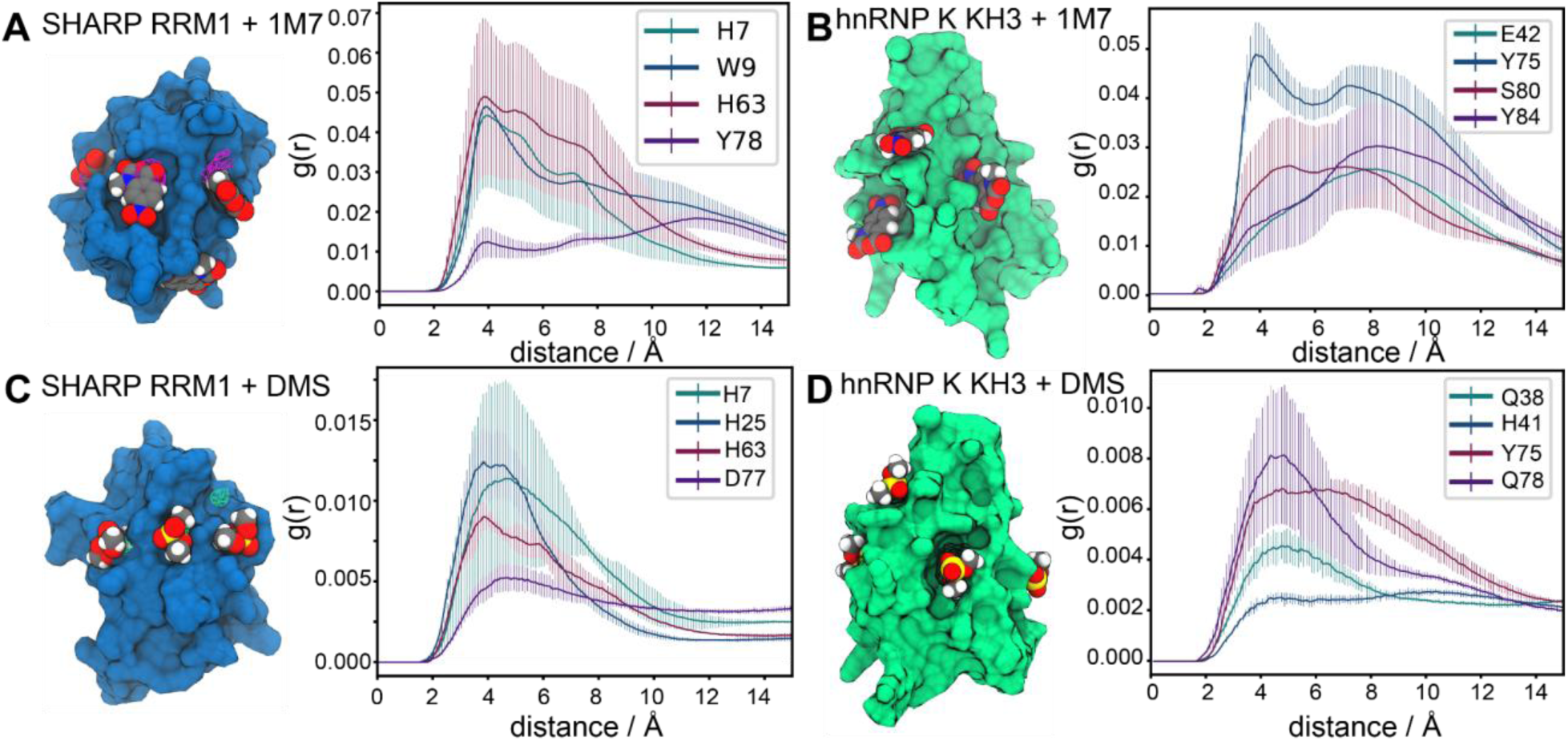
Analysis of the MD simulations of the RRM domain of SHARP with 1M7 and DMS. **A)** Simulation structure of the RRM1 domain of SHARP (blue) with 1M7, and contours representing regions of high 1M7 density (magenta). **B)** Simulation structure of the third KH domain of hnRNP K (green) with 1M7, and contours representing the regions of high 1M7 density (blue). **C)** Simulation structure of the RRM1 domain of SHARP (blue) with DMS, and contours representing regions of high DMS density (green). **D)** Simulation structure of the third KH domain of hnRNP K (green) and contours representing regions of high DMS density (blue). The plots represent the RDF of DMS or 1M7 to specific amino acids, averaged across the three simulation trials and reported with standard deviations.

DMS is smaller than 1M7, is not aromatic, and does not appear to associate with the protein surfaces with as long of a lifetime as 1M7 does in our MD simulations (**Table S1**). Instead, DMS is observed to transiently associate with polar or aliphatic residues on the protein surfaces. In the simulations of SHARP RRM1 with DMS, the DMS molecules localize near Tyr29, Asn64, and Asp77, which brings them adjacent to the potentially reactive amino acids His7, His25, and His63 (**Figure 4C**). In the simulations of the KH3 domain of hnRNP K, the DMS molecules associate with Gln38, Leu76, and occasionally Gln78, bringing them adjacent to His41 and Tyr75 (**Figure 4D**).

In summary, in both sets of simulations containing 1M7, with either SHARP or hnRNP K, the 1M7 molecules associate near potentially catalytically active amino acids with lifetimes of 50 - 100 ns (**Table S1**). Conversely, in our simulations of both proteins, the DMS molecules appear to interact with aliphatic or polar amino acids on the protein surfaces with much faster lifetimes, binding and unbinding on the order of 10 - 30 ns. Thus, the DMS molecules exhibit a more diffuse occupancy cloud around the protein than 1M7 – represented by the greater average distance of DMS probes to KH3 than 1M7 (**Figure S5**). The more transient binding interactions between DMS and the two proteins seen in our MD simulations offers a potential rationale as to why no clear signs for DMS induced methylations were observed for either protein.

### NMR spectroscopy sheds light on modification sites in hnRNP K KH3

We used NMR to experimentally validate the interaction between 1M7 and DMS with the hnRNP K KH3 protein. As a control, we first carried out a 2D ^1^H ^15^N HMQC on hnRNP K KH3 and observed good dispersion of amide resonance peaks, indicating a properly folded protein. We then titrated a 15 nucleotide long polyC RNA oligonucleotide into hnRNP K KH3 and observed line broadening and chemical shift perturbations (CSP) for several residues (**Figure 5A**). Then we recorded NH backbone spectra for hnRNP K KH3 treated with 1M7 and DMS (**Figure 5B-C**).

**Figure 5.**
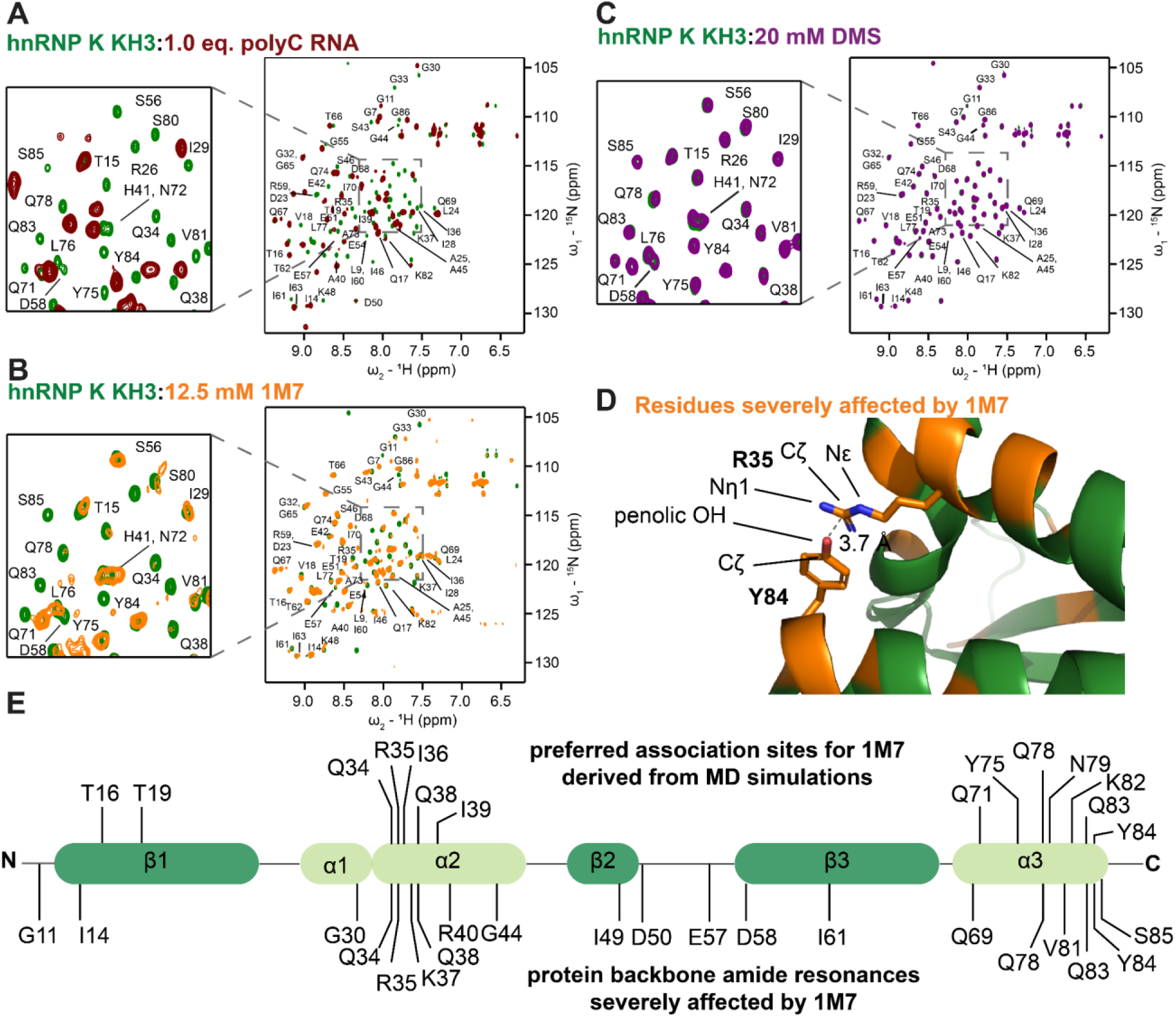
Backbone HMQC spectra recorded at 298K demonstrate binding of hnRNP K KH3 by RNA and interactions with chemical probes. Assignments refer to apo protein since CSPs and line broadening impede transfer of the assignments to bound/modified protein for many residues. Overlays show hnRNP K KH3 backbone spectrum (green) upon addition of **A)** 1.0 eq. of 15 nt polyC RNA, **B)** 12.5 mM 1M7, and **C)** 20 mM DMS. **D)** Residues severely affected by 1M7 (orange) mapped the X-ray structure of hnRNP K KH3. A stacking interaction between Arg35 and Tyr84 potentially catalyzes the reaction. The full structure is shown in **Figure S6**. **E)** Residues of hnRNP K KH3 affected by treatment with 1M7 identified by MD simulations (top) and NMR spectroscopy (bottom). At helix three and two NMR and MD results overlap nicely.

Modification of the protein with 1M7 results in substantial changes in the observed NMR spectra (**Figure 5B**). While the presence of several distinct resonances and overall good amide backbone distribution indicates a properly folded protein, almost all resonances are affected by the modification. Most backbone resonances experience some degree of line broadening or CSP; some are broadened to undetectability or escape assignment by severe chemical shift perturbations. We mapped the most severely affected resonances on the published crystal structure of hnRNP K KH3 to elucidate potential modification sites (**Figure 5D-E and S6**) (38). These are in good agreement with the association sites identified by MD simulations. Our simulations showed a preference for occupancy at helix three and its aromatic amino acid side chains. Some of the protein backbone NH resonances most affected by 1M7 are also found in this region. Additionally, we observe occupancy at helix two, wherein 1M7 molecules associate with a region including residues Q34, R35, I35, Q38, and I39. This overlaps nicely with a second cluster of NH backbone resonances affected by 1M7.

As shown, many of the residues most affected by 1M7 cluster in defined regions of the protein, suggesting a degree of systematic modification by the 1M7 chemical probe. One cluster in particular caught our attention: a multitude of amino acid residues (Gly30, Gln34, Lys37, Gln38, Val81, Gln83, Ser85) strongly affected by 1M7 are found in close proximity to Arg35 and Tyr84 (**Figure 4B**). Our simulations already suggest a preferred binding site for the 1M7 reagent near this site due to the aromatic nature of the tyrosine side chain. Interestingly, the crystal structure shows an interaction between Tyr84 and Arg35. Considering the reaction mechanism by which 1M7 modifies biomolecules, this tyrosine-arginine motif should be highly susceptible to 1M7. We speculate that the positively charged arginine side chain stabilizes a transiently deprotonated tyrosine. The resulting phenolic anion is highly nucleophilic and can attack 1M7. Catalysts promoting deprotonation have been shown to enhance the reactivity of target RNA residues for 1M7 in a similar manner (12).

What was most notable from our results is that a significant overlap exists between residues involved in the binding of polyC RNA to hnRNP K KH3, and residues affected by the 1M7 chemical probe. This suggests that such modifications can potentially impede important binding interactions (**Figure 1D, E**), thus reducing affinity. We attempted to titrate a polyC oligonucleotide into the 1M7-modified protein to study the impact on binding residues by NMR, however the extensive line-broadening made it difficult to obtain unambiguous data (**Figure S7**).

In line with our MALDI-MS data, treatment with DMS results in only minor changes of linewidths in the spectra (**Figure 5C**). Further research is needed to investigate whether DMS does indeed modify the surface of hnRNP K KH3, or a different methodology is required to identify the small changes introduced by single methyl groups.

### Reducing agents are not a solution to undesired protein modifications

We were interested in potential solutions to the undesired side reactions of chemical probing in the presence of proteins. In our study, the use of DTT in protein buffers - even if only partially - mitigates the modification of hnRNP K KH3 by 1M7 (**Figure S8**). However, the reason for this is the quenching of the chemical probing reaction by DTT (*i.e.* acetylation of DTT instead of the protein). This process is even sometimes used by design to terminate SHAPE reactions (39). Consequently, DTT does not specifically inhibit or reverse the reaction of the probing reagents with proteins but with all susceptible biomolecules, thus also impeding the desired reactivity. The same applies to all commonly used reducing agents.

### Chemical probes impede RNA binding of WDR5 protein

Considering that chemical probing experiments are typically carried out on RNAs longer than the short oligonucleotides recognized by the RBPs described in the aforementioned sections, we turned our attention to a system that could more accurately reflect the implications of protein-modification by the chemical probing reagents. We chose the interaction between the protein WD repeat-containing protein 5 (WDR5) and the long noncoding RNA (lncRNA) UMLILO, which has recently been described in literature (40).

Based on our previous results, we chose 1M7 as an exemplary SHAPE reagent. As before, we used MALDI-MS to investigate the covalent modification of WDR5 by 1M7 (**Figure 6A**). Given the substantially larger molecular weight of this protein, the resolution of the MALDI mass spectra does not permit the observation of distinct modification patterns seen for hnRNP K KH3 and SHARP RRM1 (**Figure 3**). However, an increased average molecular weight is observed for WDR5 treated with 3 mM 1M7. The amount of 1M7 was reduced because our standard procedure (18 mM) did not yield sufficient quality data (**Supporting Figure S9**). Based on our observations on hnRNP K KH3 and SHARP RRM1, we attribute this increase in molecular weight to covalent modification of the protein by 1M7. We did notice the presence of an apparent dimer peak in the treated WDR5 sample (**Figure 6A**, bottom), but further research is required to pinpoint the cause of this observation in response to treatment with chemical probes.

**Figure 6.**
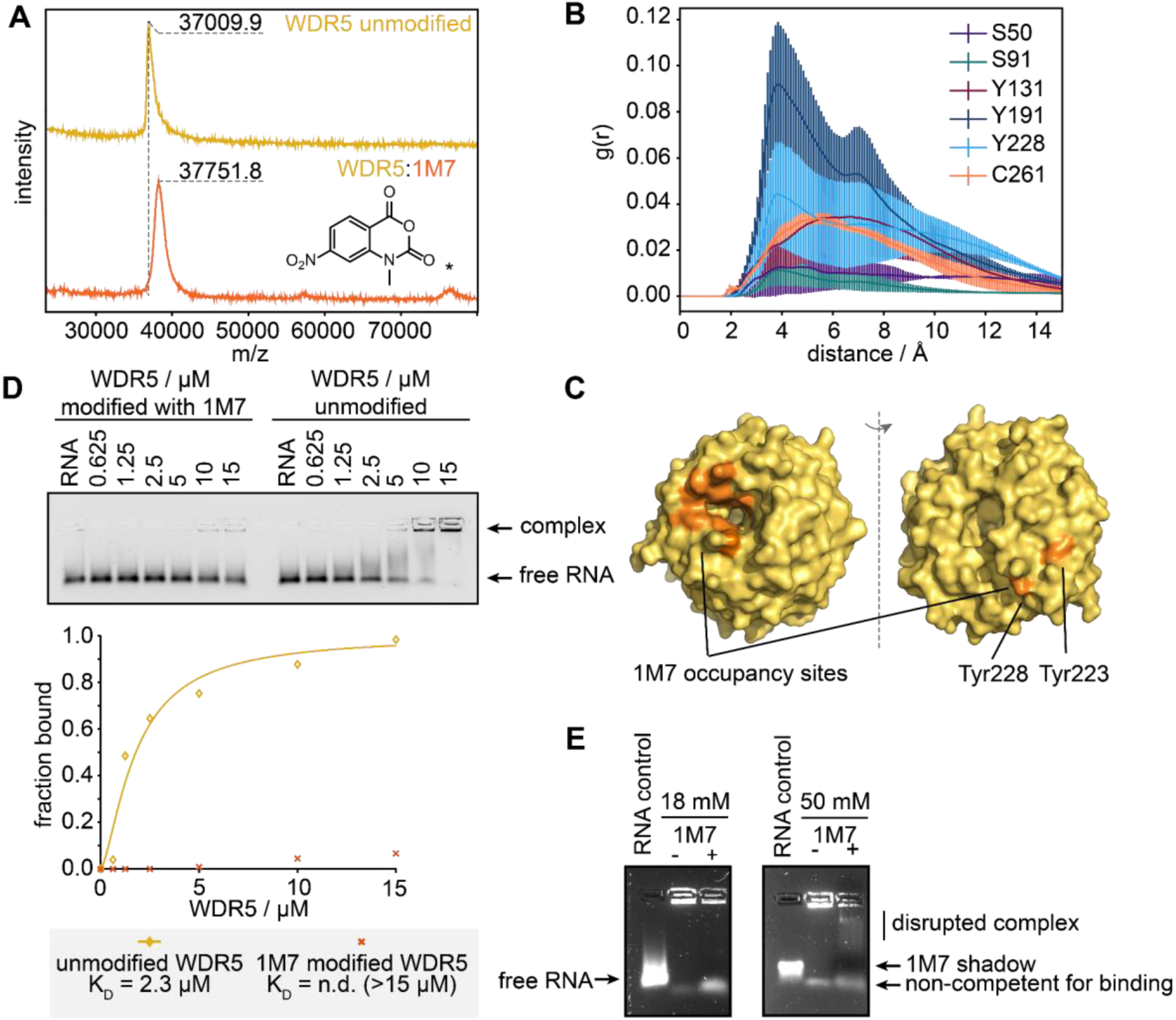
Treatment with chemical probes perturbs RNA binding of WDR5. **A)** MALDI-MS shows an increase in molecular weight of WDR5 treated with 1M7. Potential dimer formation of modified protein is highlighted with an asterisk. **B)** Radial distribution plot of 1M7 against specific residues of WDR5, averaged across three simulation trials and reported with standard deviations. **C)** Preferred 1M7 occupancy sites (orange) cluster on the structure of WDR5 (yellow). **D)** Electrophoretic mobility shift assay of WDR5 and lncRNA UMLILO. WDR5 loses its binding ability to the target RNA when treated with 1M7. A replicate of this experiment is shown in the SI (**Figure S7**). **E)** WDR5-RNA complex treated with 18 (left) and 50 mM of 1M7 results in partial disruption of the complex. The observed intensity for the complex is decreased for the treated complex at 18 mM 1M7. At 50 mM a smearing effect indicates disruption of the complex.

We then performed MD simulations of WDR5 in the presence of 1M7 to investigate sites with a propensity for 1M7 induced modifications. 1M7 molecules localize in two distinct regions with high-occupancy, preferentially interacting with aromatic surface residues of WDR5 – indicating which residues may exhibit a greater propensity for covalent modification. Specifically, we observe a strong tendency for 1M7 to localize near an area of WDR5 enriched in aromatic residues: Tyr131, Phe133, Phe149, Pro152, Tyr170, Phe219, and Tyr260 – which frequently positions 1M7 molecules near the potentially nucleophilic C240 (**Figure 6B**). Additionally, we observe a significant localization of 1M7 molecules near a known RNA binding site of WDR5, which harbors the potentially reactive residues Tyr228 and Tyr243 (41). The association of 1M7 in this RNA binding region encouraged us to investigate whether introduction of a chemical probe could inhibit the protein’s interaction with RNA – as it would in a chemical probing experiment carried out in the presence of protein.

To this end, we performed protein-RNA binding assays with WDR5 and UMLILO. Indeed, our data clearly shows how 1M7 modified WDR5 loses its binding competence for the lncRNA UMLILO. Unmodified WDR5 binds UMLILO with 2.3 μM affinity (**Figure 6D**). When treated with 1M7, however, the binding affinity is greatly reduced (**Figure 6D, S10**) to a point where the Kd could not be determined within the concentration range of the binding shift assay (> 15 μM). This demonstrates that chemical probes can introduce modifications to proteins that impede their binding to target RNAs.

Next, we tried to mimic the procedure of SHAPE experiments in the presence of proteins as closely as possible: establishing an UMLILO-WDR5 complex, followed by treatment of the complex with the 1M7 chemical probe. We combined 300 nM UMLILO RNA and 15 μM protein and incubated the complex for 20 minutes. Under these conditions, unmodified WDR5 fully saturates the RNA as shown by our EMSA data (**Figure 6D**). We treated the complex with 1M7 (or DMSO as a control) and incubated for another 20 min before native agarose gel electrophoresis. Though the complex is retained in the well, upon addition of 18 mM 1M7, the intensity corresponding to the UMLILO-WDR5 complex band is reduced. Upon increasing the concentration of 1M7 to 50 mM, this effect is exacerbated: smears corresponding to a weaker complex formed by UMLILO and WDR5 are visible on the gel (**Figure 6E**). In summary, our results demonstrate that the protein is likely subject to modifications, which occur within timescales that encompass both the on-off rate of complex formation, and the rate by which RNA is modified. This modification can inhibit its interaction with the RNA even when the complex is pre-formed. Further research is required to identify the kinetics of WDR5 complex formation and their link to modification.

## DISCUSSION

The experimental design of chemical probing against RNA-protein complexes generally involves the formation of the RNA-protein complex of interest to protect protein binding nucleotides before treatment with chemical probing reagents. Using MALDI-MS, NMR, and binding assays complemented by MD simulations, we established that RNA chemical probes are capable of selectively modifying proteins and interfering with their binding to target RNAs. We posit that this modification is possible given the life-time of the RNA-protein complex: if the life-time is sufficiently low (i.e. shorter than the half-life of the chemical probing reagent) the free protein can be modified by the chemical probing reagent, potentially causing a decrease of affinity over time, or obscuring accurate deduction of binding affinities in general. The most commonly used chemical probing reagents, with the exception of BzCN, have half-lives that are magnitudes higher than the lifetime of many RNA-protein complexes, which have been reported in a wide range between 0.1 seconds and 55 hours (42–44).

In this study, we investigated a series of chemical probes and their reactivity with proteins: 2A3, BzCN, NMIA, 1M7, and DMS. While covalent modification of the proteins by 2A3, NMIA, and 1M7 is supported by our experimental results, we did not observe strong modification patterns with BzCN or DMS. The lack of reactivity of these probes for the proteins in our study may be due to a more diffuse occupancy and a faster dissociation rate, as seen in our MD simulations with DMS, when compared to 1M7. In extension of this idea, we hypothesize that non-covalent binding of the chemical probes to the protein surface influences the preferred modification sites, but is not the primary driving force behind the chemical modification reaction – *i.e.* not all of the binding events lead to chemical modification. This also offers a rationale for why DMS-induced methylations were only observed in some of the previous studies mentioning the effect of DMS on proteins (26, 27), and highlights how a protein’s 3D structure is a critical factor for the susceptibility to chemical probing reagents. We speculate that chemical probes could potentially be used as simple means to reveal nucleophilically reactive and accessible amino acid side chains in proteins. This technique would, similar to chemical probing of RNA, involve treating a protein of interest with varying concentrations of different reagents. The introduced modifications can then be identified by a method of choice, such as high-resolution MS/MS or NMR, revealing the reactive amino acids within the protein. Similar techniques have been successfully employed to investigate properties such as hydrophobic pockets as well as structure-function relationships of proteins (45, 46).

Our UMLILO-WDR5 results demonstrate that chemical probing experiments with protein-modifying probes like 1M7 should be used with caution; they can interfere with complex formation. Furthermore, chemical probing experiments cannot be reliably used to determine binding affinities between RNA-protein complexes without further validation using orthogonal measures. Interestingly, we found a pronounced concentration dependence of the 2A3 modification rate (**Figure 3B**). Thus, the ability of a chemical probe to modify the protein relates to an intricate balance between sufficient chemical reactivity towards amino acid side chains, and a half-life long enough to access reactive sites in the protein. Keeping the concentration of the chemical probes as low as possible will also reduce the rate of undesired side reactions. We suggest that each chemical probing reagent is tested for protein side chain interactions in the future before use in RNA-protein complexes.

While not explicitly observed in this study, the possibility of non-covalent binding of chemical probing reagents and their hydrolysis products perturbing RNA binding cannot be ruled out based on our data. The potential impact of non-covalent binding largely depends on the chemical nature of the compounds. However, we expect the hydrolysis products of chemical probing reagents to be more influential than non-covalent binding in disrupting RNA-protein interactions since the supply of chemical probe will deplete over time. DMS hydrolysis yields sulfuric acid and methanol via monomethyl sulfate (47, 48). Both compounds, given sufficient buffer capacity, should not impede binding events between proteins and nucleic acids. 1M7 forms 2-(methylamino)-4-nitrobenzoic acid when reacting with water (49). This small molecule still possesses a variety of functional groups that may potentially interact with the protein, such as a carboxylic acid and amine group. Thus, we conclude that non-covalent binding of small molecule side products of a SHAPE reaction can still be relevant in some cases.

In summary, we demonstrate that chemical probes that are used to identify single-stranded and flexible nucleotides can also interact with amino acids. Many protein-RNA interactions occur over time scales from sub-seconds to hours (42). Considering that the half-lifes of chemical probing reagents are found in the same range this allows the probe to interact with the protein, potentially inhibiting its binding (39). Critically, the reaction of a chemical probe with amino acid residues that are known to facilitate interactions with RNA (e.g. tyrosine) can diminish the binding interaction. On a similar note, RNA-binding molecules (e.g. therapeutic agents) can also be modified by a chemical probe, and its interaction with its RNA target inhibited. As has been observed in multiple studies, even at sites of ligand binding, nucleotides can still possess chemical probe reactivity (50–53). As a silver lining, RNA-modifying probes could serve as a useful tool to identify surface residues of proteins, and could potentially be used as an alternative to introducing mutations to amino acids on the RNA-binding interface. Future studies will investigate the factors that govern the reactivity of protein amino acids with chemical probing reagents, and the mechanisms by which they perturb RNA-protein interactions.

## EXPERIMENTAL PROCEDURES

### Protein Expression

Plasmids encoding for SHARP RRM1 (9.5 kDa) and the KH3 domain of hnRNP K (9.0 kDa) (each preceded with an N-terminal 6x histidine tag (his-tag) and a tobacco etch virus (TEV) cleavage site) were transformed into BL21 DE3 *E. coli* chemically competent cells and expressed in LB or M9 minimal media solution supplemented with ^15^N NH4Cl. Cells were grown at 37°C; upon reaching an optical density (OD600) of 0.9, they were induced to express protein with 0.5 M isopropyl ß-D-1-thiogalactopyranoside (IPTG) at 18°C. Cells were lysed by sonication and purified using immobilized metal (nickel) affinity chromatography (IMAC). Briefly, the protein was washed in a buffer containing 50 mM Tris pH 8, 300 mM NaCl, 5 mM beta-mercaptoethanol, and 10 mM imidazole. An on-column cleavage of the his-tag was performed by adding 10 mg/mL of TEV protease. The flow through was assessed by sodium dodecyl-sulfate (SDS) polyacrylamide gel electrophoresis (PAGE) followed by size exclusion chromatography (HiLoad 16/60 Superdex 75) in a buffer containing 150 mM NaCl, 5 mM dithiothreitol (DTT), and 25 mM sodium phosphate, pH 6.5. Purified fractions were assessed by SDS PAGE, followed by flash freezing in liquid nitrogen and storage at -80°C until further use.

Plasmid encoding for WDR5 (36.6 kDa) was obtained from addgene (Addgene plasmid # 59969) and ligated into pETM11 with an N-terminal 6x-his tag and TEV cleavage site. The plasmid was transformed into BL21 DE3 *E. coli* competent cells. LB was inoculated with an overnight culture and incubated at 37°C until OD600 reached 0.8. Expression was induced with 1 mM IPTG at 18°C overnight. Cells were lysed by sonication in a buffer containing 25 mM Tris, pH 7.5, 300 mM NaCl, and 5 mM DTT then purified by IMAC, eluted with 300 mM imidazole followed by dialysis and cleavage with TEV protease. Protein was further purified by size exclusion chromatography on a Superdex S75 column in a buffer containing 25 mM sodium phosphate, pH 7.5, 150 mM NaCl, and 5 mM DTT. Fractions were analyzed by SDS-PAGE, flash-frozen, and stored at -80°C.

### RNA synthesis

The DNA template encoding for UMLILO was amplified by PCR from a plasmid with the exonic sequence (a gift from Dr. Musa Mhlanga) using a forward primer containing the T7 promoter sequence. The template DNA was cleaned using AmpureXP bead-based reagent. The transcription reaction consisted of 10-20 ng/µL template DNA, 8 mM of each rNTP, 10% polyethylene glycol 8000, 20 mM MgCl2, 1X transcription buffer (5 mM Tris, pH8, 5 mM spermidine, 10 mM DTT), and in-house T7. RNA was in vitro transcribed at 37°C for 3 hours then purified by high-performance liquid chromatography (HPLC) using 12.5 mM Tris HCl and 6M urea and eluted with a gradient of 0-500 mM sodium perchlorate.

### Protein Modification

Protein samples with a concentration of 200 µM of the respective protein in a buffer (25 mM sodium phosphate, 150 mM NaCl, pH 6.5) were prepared. Stock solutions of 1M7 (a gift from Michael Sattler), NMIA, BzCN, and 2A3 in anhydrous DMSO were added for a final concentration of 18 mM (12.5 mM for NMR studies of KH3 and 3 mM for MALDI-MS of WDR5) 1M7 in the reaction mixture. For DMS, 40 µL DMS were dissolved in 90 µL of anhydrous ethanol and diluted with 870 µL of water. Aliquots of this solution were then added to 200 µM protein solutions for a final concentration of 20 mM DMS. The mixtures were incubated for 15 minutes at room temperature prior to use.

### NMR Analysis

To eliminate contaminations by side products of the modification reactions, the modified proteins were loaded onto a 3 kDa cutoff spin column. Four subsequent washings with fresh NMR buffer (25 mM sodium phosphate, 150 mM NaCl, pH 6.5) restored the initial solute composition. After recovery, the samples were supplemented with 10% D2O and transferred into 3 mm NMR tubes. Sample concentrations were in the range of 100-120 µM (absorption of 1M7 inhibits concentration determination of 1M7 modified proteins by UV). NMR experiments were conducted on a Bruker 800 MHz Avance NEO equipped with a cryo-probe at 298 K. The 2D ^1^H-^15^N correlation spectra were acquired using a SOFAST-HMQC pulse sequence (54). Spectra were processed in Topspin 3.6 and analyzed with CCPN2.5.3 (55). Protein backbone resonance assignments for hnRNP K KH3 have been published elsewhere and could be transferred to our spectra for most backbone resonances without complications (56, 57). For the RNA titration experiments 5’-(CCC)5-3’ RNA and DNA oligonucleotides were supplied by Integrated DNA Technologies.

### MALDI Mass Spectrometry

All mass spectrometric experiments were conducted on a Bruker Autoflex maX MALDI-TOF instrument. All spectra were acquired in positive mode. Aliquots of the (modified) protein samples were diluted to <6 µM with water and combined with α-Cyano-4-Hydroxycinnamic acid for hnRNP K KH3 and SHARP RRM1 and 2, 5-dihydroxybenzoic acid for WDR5 using the dried droplet technique. WDR5 samples were additionally desalted by washing with water on a 3 kDa cutoff spin column. This step was omitted for hnRNP K KH3 and SHARP RRM1 since we found no beneficial effect on spectral quality for these proteins. Mass spectra were recorded for SHARP RRM 1 modified with 1M7 and DMS; hnRNP K KH3 modified by 1M7, DMS, NMIA, BzCN, and 2A3; WDR5 modified by 1M7 as well as the unmodified proteins.

### Molecular Dynamics Simulations

1M7 and DMS were separately docked to the WDR5, the RRM1 domain of SHARP, and the KH3 domain of hnRNP K. A structure for the RRM1 domain of SHARP was modeled using *AlphaFold2* (58) . The WDR5 protein, and the KH3 domain of hnRNP K, were taken from their respective X-ray structures (PDB IDs 8G3C and 1ZZK, respectively) (38, 59); the GB1 solubility tag was removed from the KH3 domain. Structures for 1M7 and DMS were procured from the CSD (60). Parameters for 1M7 and DMS were obtained using GAFF and Antechamber (61). The 1M7 and DMS molecules were docked to the RRM1 domain of SHARP using the flexible docking algorithm of UCSF’s *Dock6* (*62*). For each ligand, the top four ranked poses were used.

The six systems were then prepared for simulation: (i) hnRNP K KH3 + DMS, (ii) hnRNP K KH3 + 1M7, (iii) SHARP RRM1 + DMS, (iv) SHARP RRM1 + 1M7, (v) WDR5 + DMS, (vi) WDR5 + 1M7. The *tleap* module of *Amber22* (*63*) was used to solvate the system in a truncated octahedron box of 9, 833 *TIP3P* (*64*) water molecules before neutralizing the total charge with Na+ ions; a 50 mM NaCl buffer was then added. The total system size consisted of 30, 901 atoms. The protein was simulated using the *ff14SB* (*65*) force field, and the ions were modeled with the Joung-Cheatham monovalent ion set in *TIP3P* (*66*). The 1M7 and DMS molecules were modeled using *GAFF* (*61*). The masses of non-water hydrogen atoms were repartitioned to permit a 4 fs timestep in production runs (67). A distance cutoff of 10 Å was applied to nonbonded interactions, with long-range electrostatics calculated using Particle Mesh Ewald (68). A Langevin thermostat (69) was used to maintain the temperature with a collision frequency of 1 ps^-1^. Simulations were carried out using the GPU variant of the *pmemd* module of *Amber22*.

The systems were relaxed identically using a 10-step equilibration protocol designed to prepare the systems for simulation conditions. The first step included 1, 000 steps of steepest descent minimization, followed by 9, 000 steps of conjugate gradient minimization with only water molecules and hydrogen atoms unrestrained and 100.0 kcal/(mol*Å^2^) Cartesian positional restraints on the rest of the system. The second step involved heating from 100 K to 298.15 K over 1 ns at constant NVT, again with all atoms except hydrogen atoms and water molecules restrained with a 100 kcal/(mol*Å^2^) force constant, before maintaining the temperature at 298.15 K for an additional 4 ns. The third step was 1 ns MD simulation at constant NPT with 100 kcal/(mol*Å^2^) positional restraints on all atoms except hydrogen and water molecules. The fourth step was 1 ns MD simulation at constant NVT with 100 kcal/(mol*Å^2^) restraints on all atoms except hydrogen and water molecules. The fifth step was 1, 000 steps of conjugate gradient minimization with only the protein backbone atoms (Cα, C, N) restrained with a 10 kcal/(mol*Å^2^) force constant. The sixth step was 1 ns MD simulation at constant NPT with 10 kcal/(mol*Å^2^) restraints on the protein backbone, followed by another 1 ns MD at constant NPT with 1 kcal/(mol*Å^2^) restraints on the protein backbone, then another 1 ns with 0.1 kcal/(mol*Å^2^) restraints, followed by a final 10 ns of unrestrained MD at constant NPT. The four 1M7 and DMS molecules in each system were unrestrained during the final four equilibration steps. The final coordinates and velocities from the last relaxation step were used to seed triplicate independent MD simulations at 298.15 K for each system.

Analysis was carried out using the *cpptraj* (*70*) module of *Amber22*. The AlphaFold2 model was used as the reference structure for SHARP RRM1, the first model of the NMR structure was used as the reference structure for hnRNP K KH3, and the X-ray structure was used as the reference for WDR5 (38, 40). The RMSF calculations were performed by fitting the non-terminal backbone atoms of the simulated structures to the reference structure before calculating the per-residue RMSF using all non-hydrogen atoms. The RDFs of 1M7 and DMS were calculated against particular amino acids on each protein to monitor which residues DMS and 1M7 interact with the most during the simulations. Structures were visualized using VMD (71).

### Electrophoretic mobility shift assays

EMSAs were conducted in a buffer consisting of 25 mM sodium phosphate pH 6.5 and 150 mM sodium chloride. 300 nM UMLILO RNA were combined with dilutions of WDR5 (unmodified or modified with 18 mM 1M7) for final concentrations of 0, 0.625, 1.25, 2.50, 5.00, 10.0, and 15.0 μM in 10 μl. The samples were incubated at room temperature for 20 minutes before adding 3 μl 30% glycerol and loading on a 1% native agarose gel stained with SYBR safe. Gels were run at 60V for 40 minutes. Gels were using ImageJ (72). The decay of the free RNA band was quantified for unmodified WDR5 and curves fitted to the Hill equation using MatLab. For modified WDR5 both complex (remaining in the well) and free RNA were quantified to determine fraction bound in order to avoid interfering of 1M7 absorbance.

## DATA AVAILABILITY

All data are included in the article or available from the corresponding author A.N.J.

## SUPPORTING INFORMATION

This article contains supporting information.

## ACKNOWLEDGEMENTS

The authors would like to thank Emma Gogarnoiu for assistance with figures, Alex Nazzaro for assistance with MALDI experiments, and members of the Jones lab for thoughtful comments and suggestions.

