## Supporting Information for "The impact of RNA chemical probing reagents on RNA binding proteins"

† Joint Authors

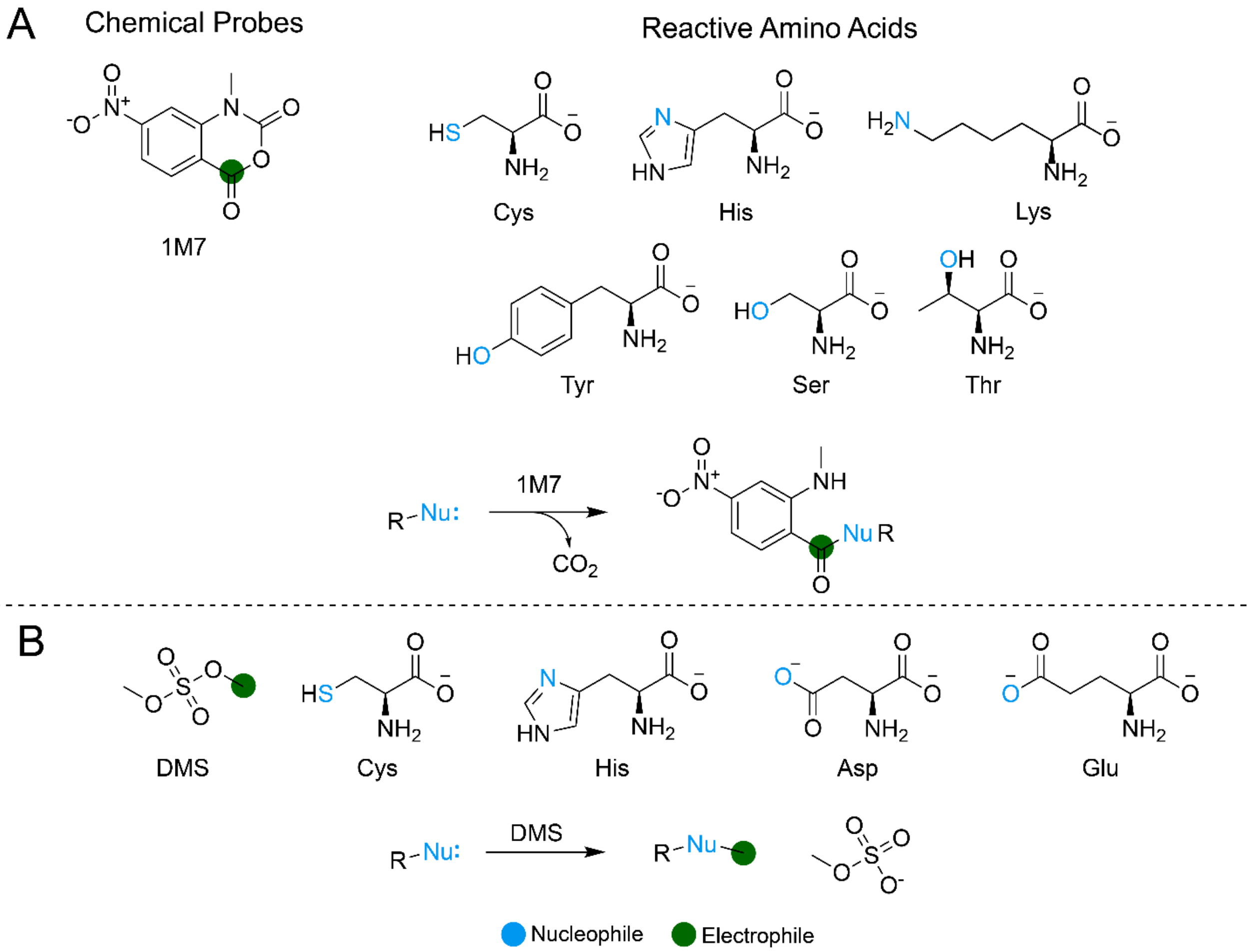


**Supporting Figure 1.** Nucleophilic amino acid side chains that react with RNA-modifying probes. **A)** Nucleophilic atoms of Cys, His, Tyr, Ser, Thr, and Lys residues with 1M7. **B)** Nucleophilic atoms of Cys, His, Asp, and Glu residues with DMS.

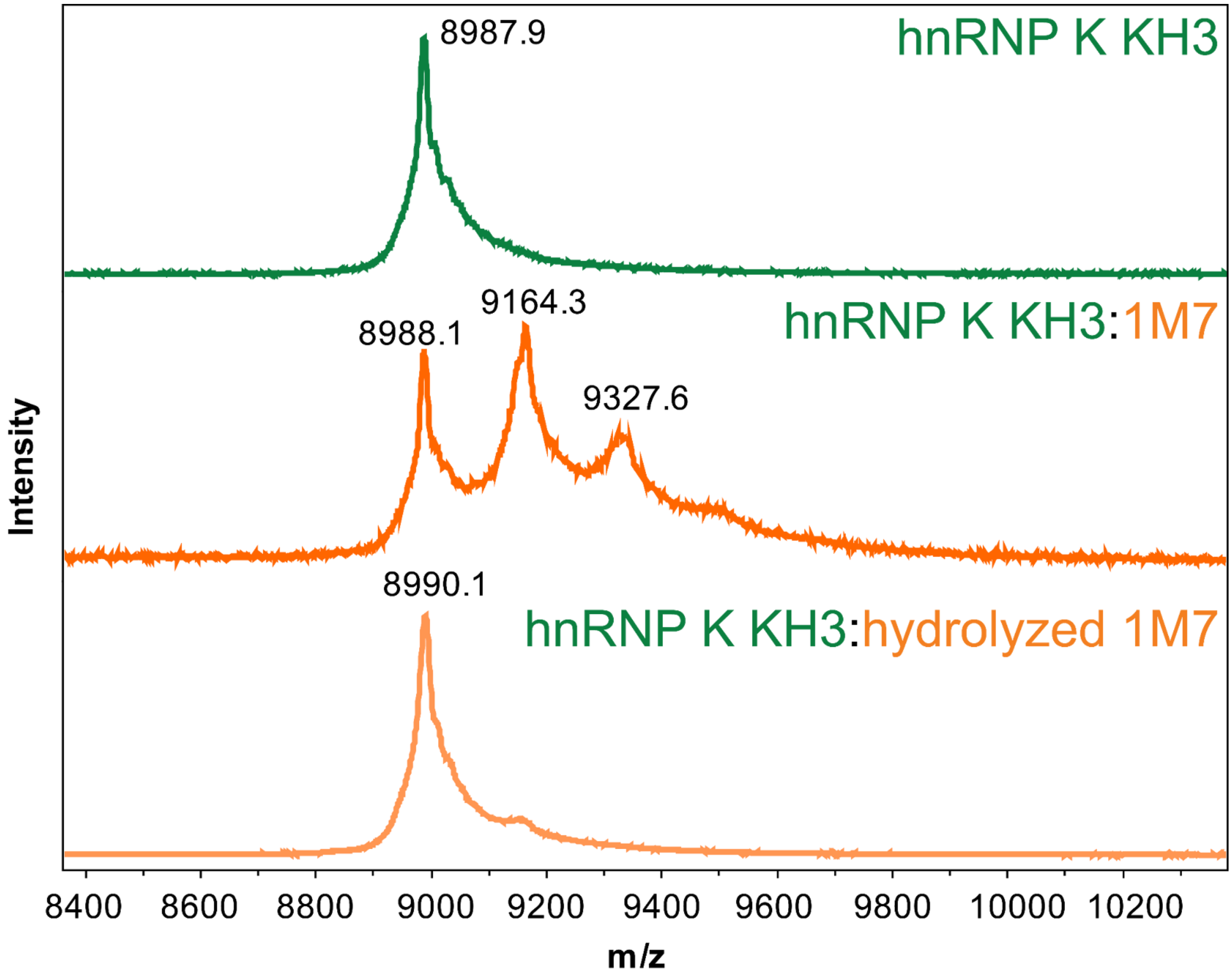


**Supporting Figure 2.** MALDI-TOF spectra showing the effects of active 1M7 (middle) and hydrolyzed 1M7 (bottom) reagents on the molecular weight of hnRNP K KH3 (top).

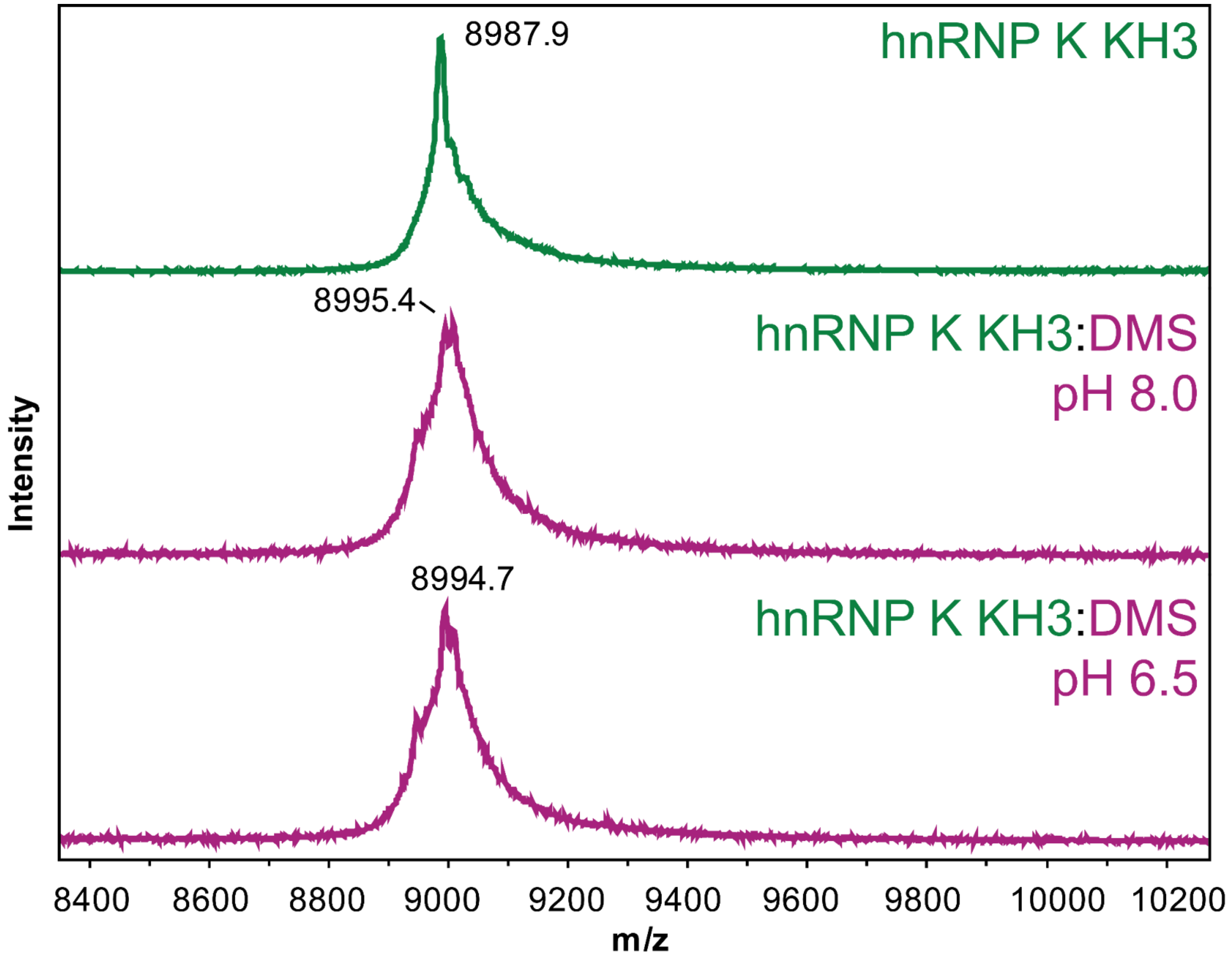


**Supporting Figure 3**. MALDI-TOF data for hnRNP K KH3 unmodified (top) and treated wit 20 mM DMS at pH 8.0 (middle) and pH 6.5 (bottom).

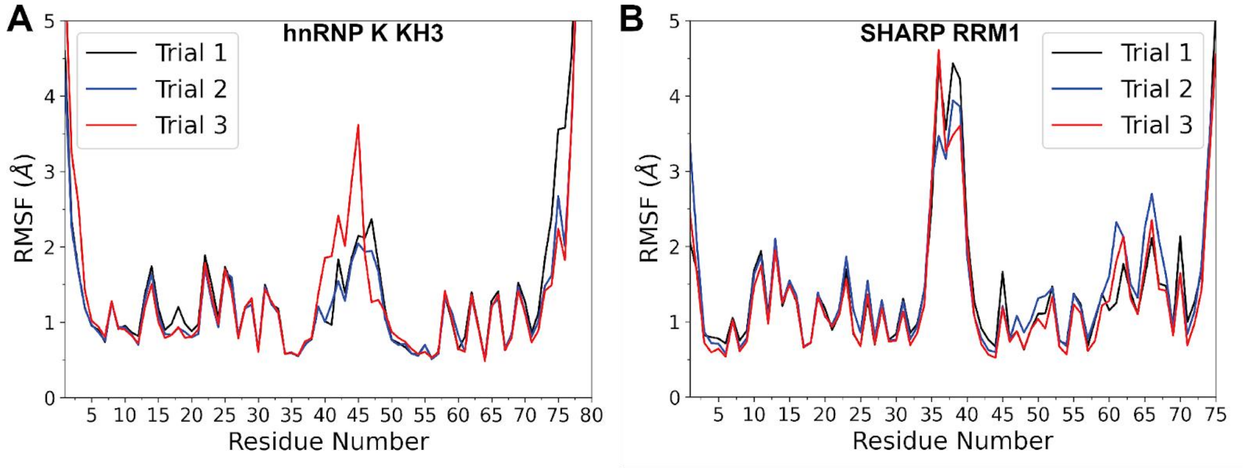


**Supporting Figure 4**. RMSF of non-hydrogen atoms in the side chains of **A)** the KH3 domain of hnRNP K, and **B)** the RRM1 domain of SHARP. The different colored lines correspond to individual simulation trials, with each system simulated for 1 μs in triplicate.

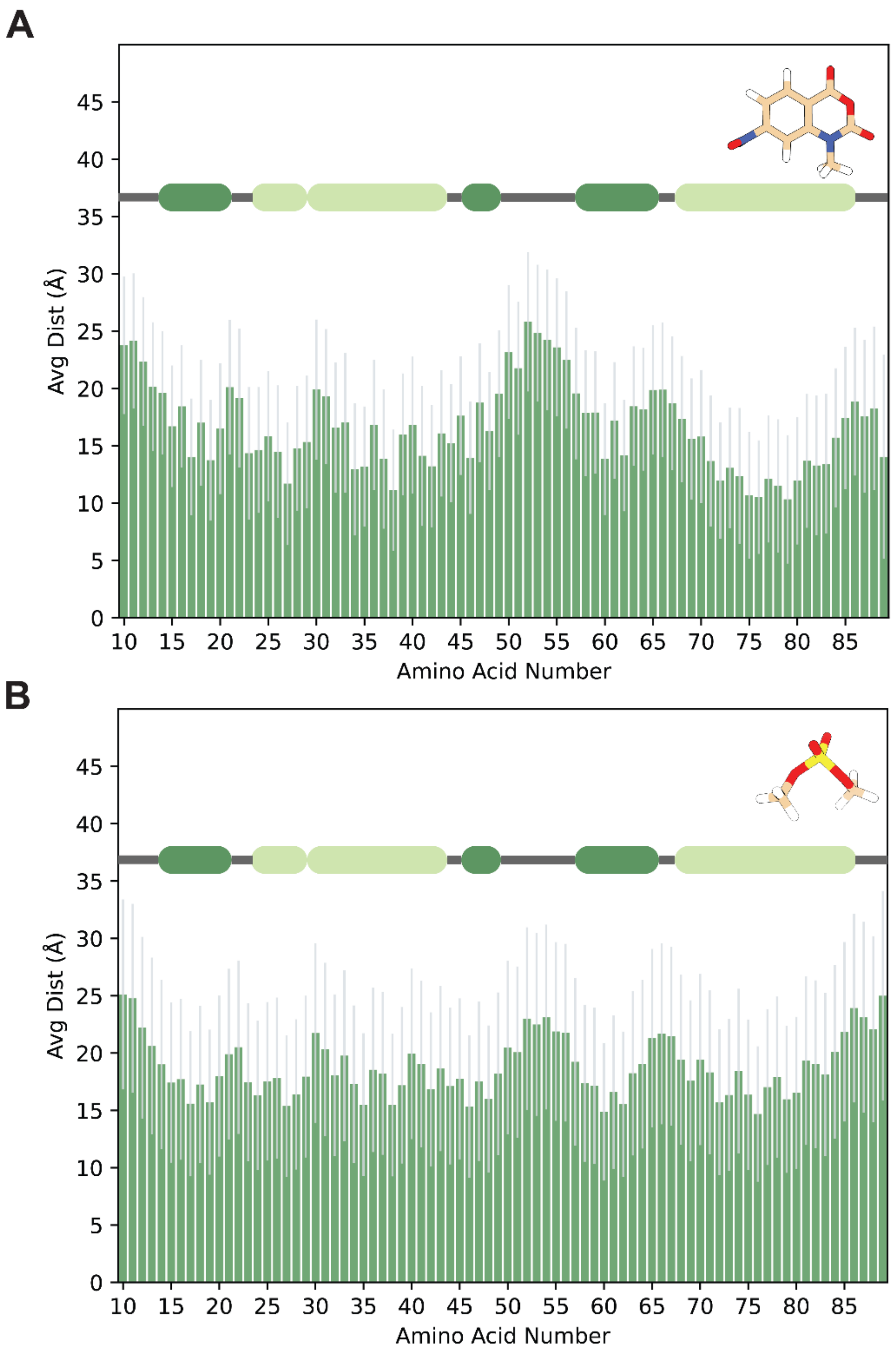


**Supporting Figure 5**. Average distance of (A) 1M7 and (B) DMS to each residue of hnRNP K KH3 throughout triplicate MD simulations. The sausage plots are colored according to secondary structure, with dark green regions representing ß-sheets, and the light green regions representing α-helices.

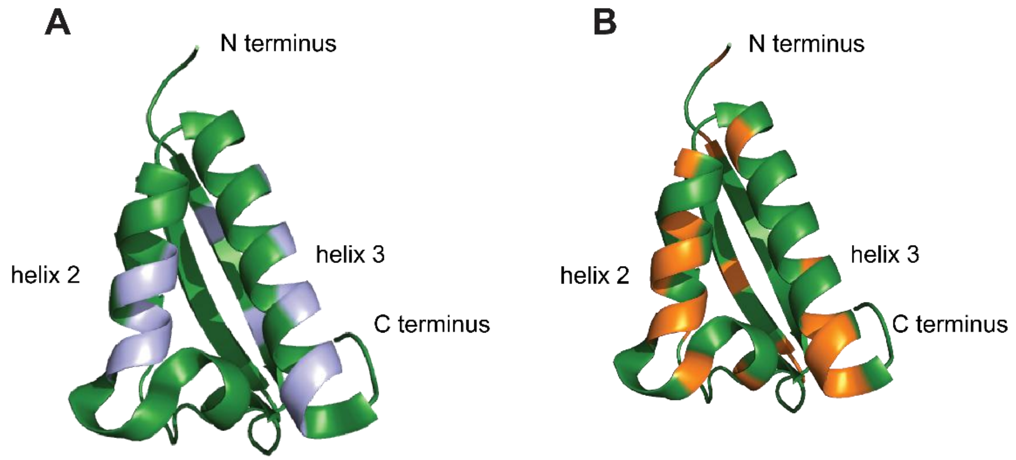


**Supporting Figure 6.** Comparison between results from MD simulations and NMR spectroscopy. **A)** Preferred association sites for 1M7 mapped on the published X-ray structure (1). **B)** Residues of hnRNP K KH3 severely affected by treatment with 1M7 chemical probing reagent mapped onto the same structure.

| | Average Binding Lifetime KH III (ns) | Average Binding Lifetime RRM1 (ns) |
|---|---|---|
| 1M7 | 86.3 ± 24.8 | 64.7 ± 16.2 |
| DMS | 28.9 ± 6.3 | 14.2 ± 5.2 |

**Supporting Table 1**. Average lifetimes of 1M7 and DMS binding events to the KH3 domain of hnRNP K and the RRM1 domain of SHARP, reported with standard deviations.

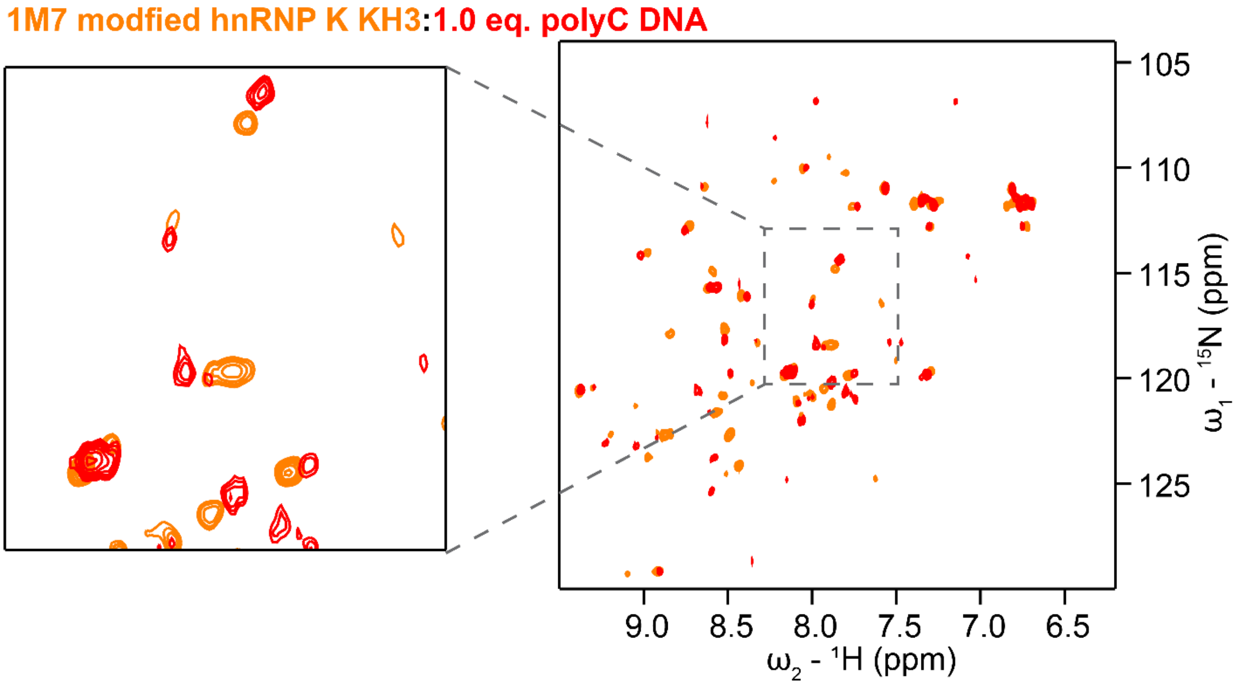


**Supporting Figure 7.** Titration of a 15 nt polyC DNA to 1M7-modified hnRNP K KH3 observed by backbone NH HMQC. Treatment with 1M7 causes line broadening and shifts leading to a complex apo spectrum (orange). Shifts can be observed upon addition of DNA (red) but due to a lack of assignments and line broadening no clear conclusions can be drawn from the spectra.

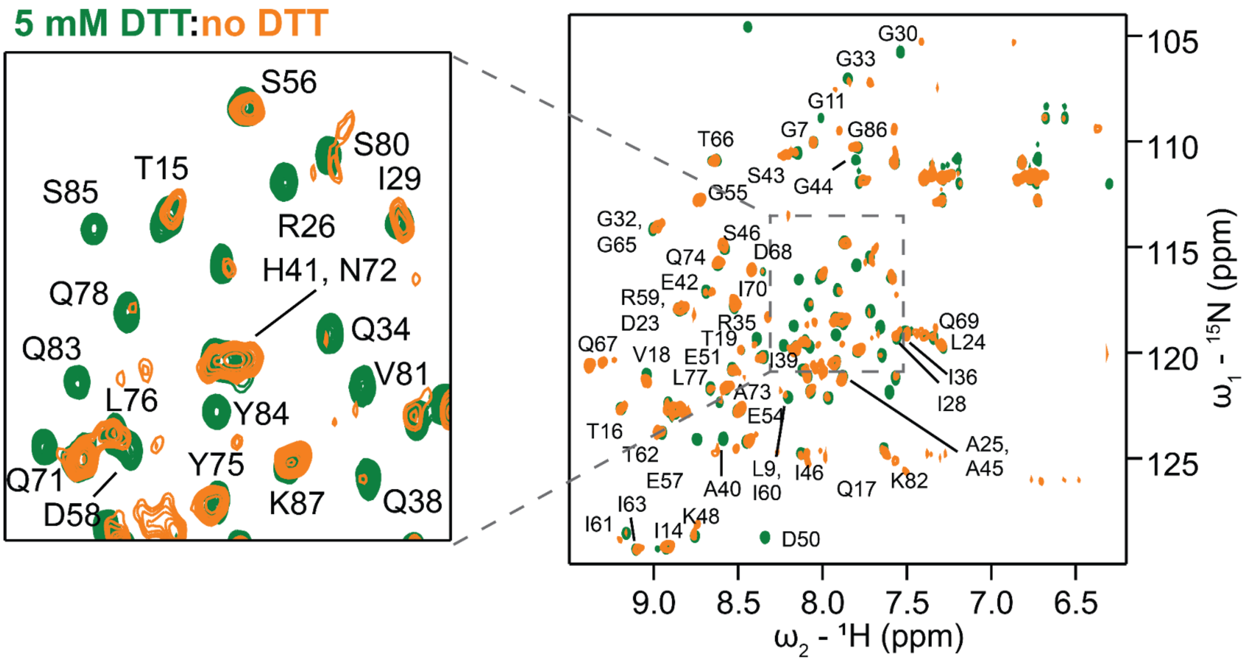


**Supporting Figure 8.** Mitigation of 1M7 modifications by DTT. The backbone NH HMQC spectrum of hnRNP K KH3 treated with 12.5 mM 1M7 in the presence of 5 mM DTT (green) and in the absence of DTT (orange).

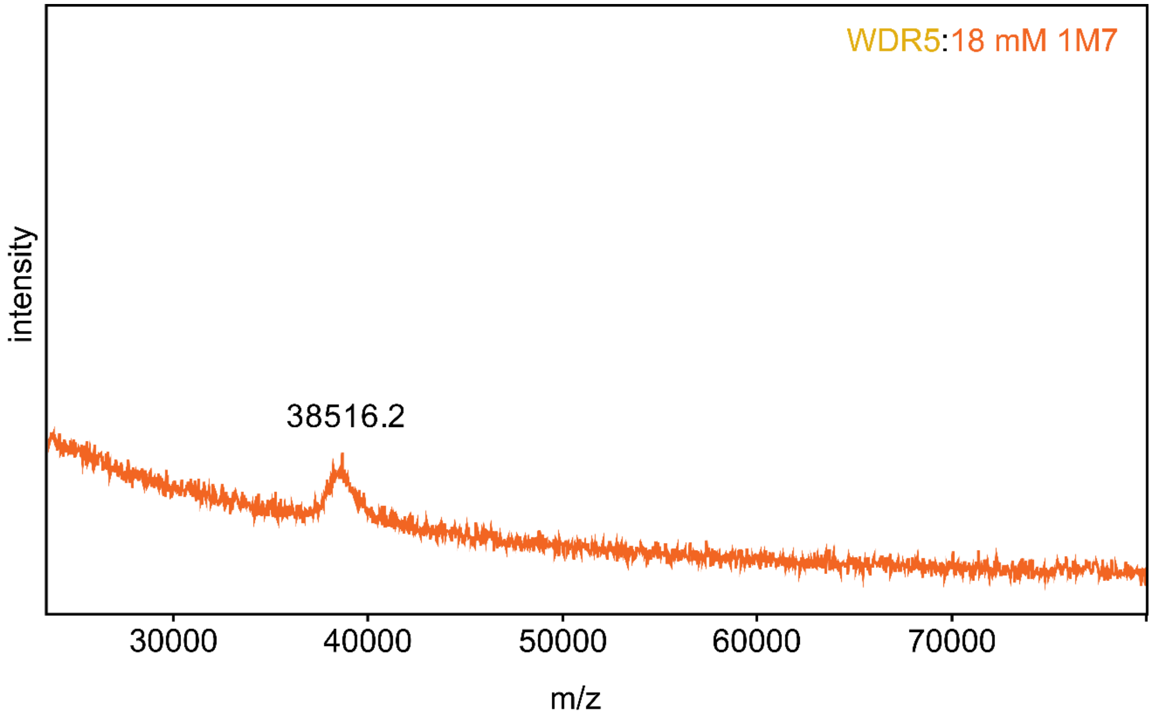

**Supporting Figure 9.** MALDI mass spectrum of WDR5 treated with 18 mM 1M7. Under these conditions the spectrum suffers from poor signal-to-noise and resolution.

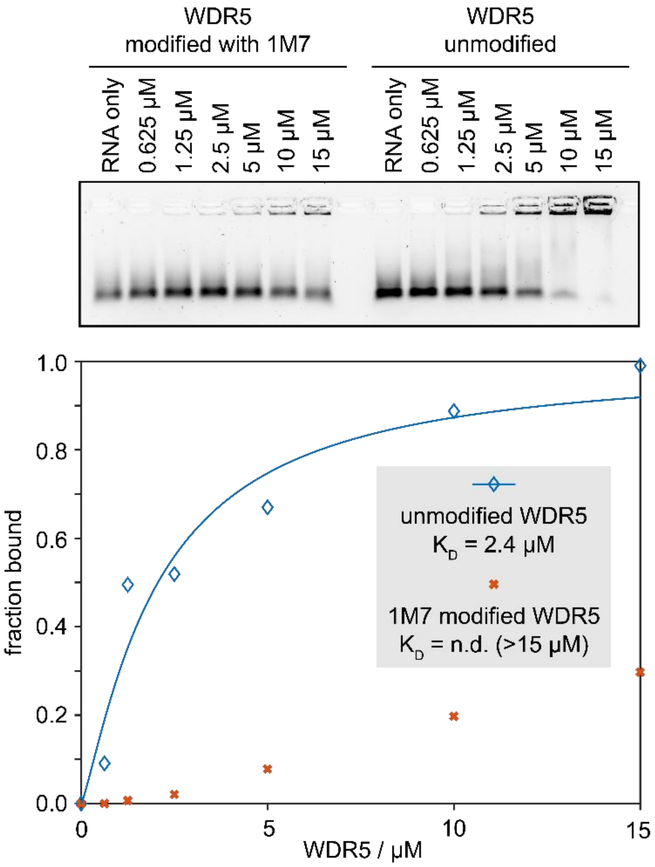


**Supporting Figure S10.** Replicate for the electrophoretic mobility shift assay shown in main text (**Figure 6D**).
